# From Lab Notes to Linked Data: MeSyTo for Ontology-Driven Metadata in Toxicological Omics

**DOI:** 10.64898/2026.08.14.736375

**Authors:** Marina Pozhidaeva, Stephan Schreiber, Kristin Schubert, Wibke Busch, Jörg Hackermüller, Sebastian Canzler

## Abstract

Toxicological omics studies require comprehensive metadata to support reproducibility, inter-operability, and regulatory reuse. However, metadata requirements differ across public repositories, reporting frameworks, and laboratory workflows, resulting in inconsistent annotation and limited data integration. To address this challenge, we developed MeSyTo (Metadata for Systems Toxicology), an ontology-driven framework for harmonizing metadata across toxicological omics. Metadata concepts from public repositories, the OECD Omics Reporting Framework (OORF), community standards, and institutional workflows were semantically aligned and implemented as the MeSyTo Metadata Model (MMM). The MMM serves as the basis for the automatic generation of SHACL validation shapes and framework-specific metadata profiles, while curated value sets are represented as SKOS controlled vocabularies to support metadata collection and validation. The current implementation comprises 105 ontology classes and 527 data properties and supports transcriptomics, proteomics, and metabolomics. A prototype web application demonstrates ontology-driven metadata collection with integrated semantic validation and ontology-based term resolution. The ontology, validation shapes, controlled vocabularies, generation scripts, and software are publicly available as open-source resources. MeSyTo provides a reusable semantic foundation for harmonized, machine-actionable metadata and facilitates repository submission, regulatory reporting, and interoperable data exchange across toxicological omics studies.

## 1 Introduction

Omics techniques such as transcriptomics, proteomics, and metabolomics are increasingly applied in systems toxicology to identify mechanisms of toxicity, discover biomarkers, and improve predictive models (Joseph, 2017; Canzler *et al*., 2020; Olesti *et al*., 2021; Madeira and Costa, 2021). For example, by enabling the integration of molecular-level observations into mechanistic frameworks such as Adverse Outcome Pathways (AOPs) (Ankley *et al*., 2010) and the establishment of transcriptomic points of departure (Reardon *et al*., 2023), omics experiments are expected to play a key role in advancing next-generation risk assessment strategies (Buesen *et al*., 2017; Brockmeier *et al*., 2017). Despite this potential, large-scale exploitation of omics in toxicology remains limited by insufficient standardization of metadata describing experimental design, materials, protocols, and data processing. Inconsistencies and gaps in metadata annotation across studies and repositories complicate data reuse, integration, and meta-analysis, and significantly hinder regulatory acceptance. Toxicological omics studies are inherently diverse in biological models, exposure scenarios, and analytical platforms, resulting in heterogeneous reporting practices and often incomplete contextual documentation (Sauer *et al*., 2017; Harrill *et al*., 2021).

In response to these challenges, several international initiatives have emerged to establish standardized metadata and reporting practices. Notably, the OECD Extended Advisory Group on Molecular Screening and Toxicogenomics (EAGMST) developed the OECD Omics Reporting Framework (OORF), a modular, integrated reporting framework designed to support the regulatory application of transcriptomics and metabolomics data (Harrill *et al*., 2021). While OORF represents the most comprehensive structural effort to date in harmonizing omics reporting within toxicology, its adoption by public repositories has been limited. Repositories such as National Center for Biotechnology Information (NCBI) Gene Expression Omnibus (GEO)^1^, European Bioinformatics Institute (EBI) BioStudies^2^, and PRIDE^3^ typically rely on existing standards and metadata checklists tailored to specific omics data types, such as Minimum Information about a high-throughput Nucleotide Sequencing Experiment (MINSEQE) (Brazma *et al*., 2012), Minimum Information About a Proteomics Experiment (MIAPE) (Taylor *et al*., 2007), and MSI (Sumner *et al*., 2007). However, these target repositories and standards neither ensure full interoperability across omics domains nor fully address the specific needs of toxicological applications, and enforcement within repositories is often limited.

For instance, while NCBI GEO provides template-based submission formats and automated validation for selected fields, much of its descriptive metadata is still captured as free text. In contrast, the OORF is currently distributed as a narrative reporting framework without a machine-actionable schema. Although EBI BioStudies supports ontology-backed fields for selected metadata elements, comprehensive semantic alignment across repositories, regulatory frameworks, and local laboratory documentation remains limited.

To enhance the reuse and machine-actionability of data, the FAIR Data Principles were introduced, promoting Findability, Accessibility, Interoperability, and Reusability of digital assets (Wilkinson *et al*., 2016). Several tools have emerged to support the creation of high-quality, FAIR-compliant metadata. For example, the FAIR Data Station (FAIR-DS) enables structured, ontology-backed metadata entry, with built-in validation of formats, restricted values, and term usage (Nijsse *et al*., 2023). It also offers flexibility through customizable templates and local configurations. However, it currently lacks features such as synonym resolution and cross-template harmonization of metadata fields.

In summary, while existing standards and tools provide valuable guidance toward transparent and reproducible reporting, they do not yet provide a harmonized, ontology-backed, and machine-actionable framework that connects repository requirements, regulatory reporting expectations, and institutional laboratory practices for toxicological omics.

To address this gap, we developed *MeSyTo* (Metadata Harmonization in Systems Toxicology), a harmonized metadata framework specifically tailored to the requirements of omics experiments in toxicology by integrating elements from OORF, GEO, EBI BioStudies, PRIDE, and internally established laboratory practices. We captured key aspects of typical workflows, such as study design, biological materials, exposure information, sample processing, analytical methods, and data analysis, balancing research and regulatory necessities with practical usability. We further implemented an ontology-driven representation of the metadata schema using established ontologies such as NCI Thesaurus (NCIt), Experimental Factor Ontology (EFO), and Ontology for Biomedical Investigations (OBI) to provide semantic interoperability and formal structure. This work therefore provides a comprehensive and extensible foundation for metadata management in toxicology. The framework supports consistent annotation across experimental and computational pipelines, promotes regulatory transparency, and contributes to ongoing efforts to establish interoperable infrastructures for systems toxicology and risk assessment.

## 2 Metadata Harmonization

In this section, we outline our metadata harmonization strategy across repositories and standards, encompassing both metadata field names (keys) and their corresponding values. This serves as the semantic foundation for ontology-based validation and integration, described in the following section. While the framework is designed to be broadly applicable across omics types, it is explicitly focused on toxicological and regulatory-relevant experiments, which introduce specific metadata requirements not typically addressed in general-purpose standards.

To enable the generation of interoperable, machine-actionable metadata for toxicological omics studies, we propose a multi-step approach that integrates existing standards, ontology mappings, and technical validation mechanisms (Figure 1).

**Figure 1:**
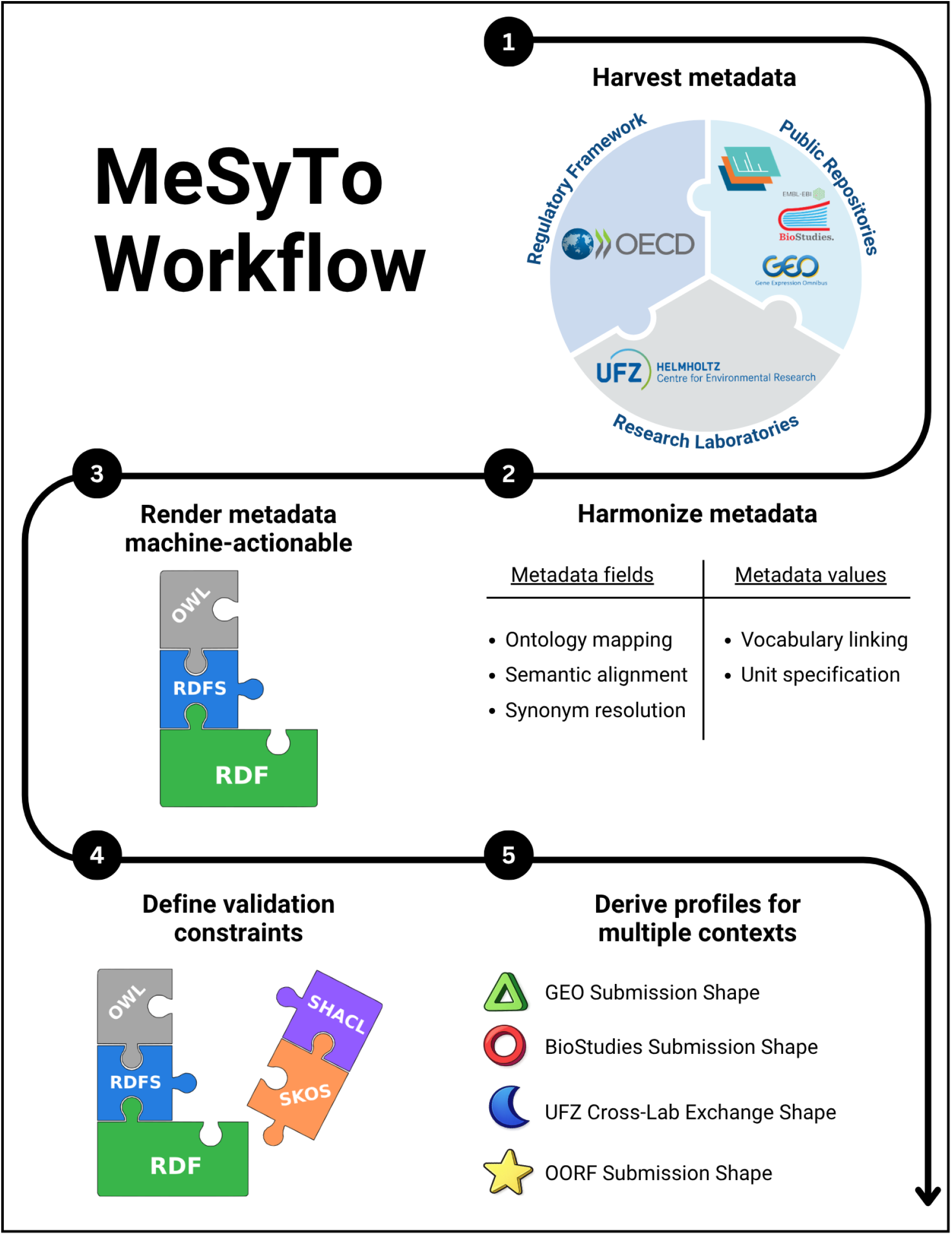
MeSyTo harmonization workflow. Overview of the MeSyTo workflow for ontology-driven metadata harmonization in toxicological omics. (1) Metadata concepts are collected from regulatory frameworks, public repositories, and laboratory workflows. (2) Metadata fields and values are harmonized through semantic alignment, ontology mapping, synonym resolution, vocabulary linking, and unit specification, and (3) represented in Resource Description Framework (RDF) and Web Ontology Language (OWL). Shapes Constraint Language (SHACL) constraints and Simple Knowledge Organization System (SKOS) vocabularies are then used to define (4) validation rules and to derive (5) profile-specific shapes for different target contexts, such as repository submission, regulatory reporting, and institutional metadata exchange.

### 2.1 Metadata Standards Landscape

Metadata standards for omics data are shaped by the distinct needs of different research communities, repositories, and regulatory frameworks. In this section, we briefly review key sources that informed our harmonization approach, with particular attention to their scope, structure, and relevance for toxicology.

#### Transcriptomics

The MINSEQE standard defines the essential metadata required to interpret and reuse transcriptomics data, including sample attributes, experimental design, and data processing steps (Brazma *et al*., 2012). The most widely used repositories for transcriptomics data, the GEO and BioStudies, recommend MINSEQE compliance, but differ in their implementation and enforcement. GEO accepts submissions using official Excel metadata templates and encourages users to provide MINSEQE-compliant information. For high-throughput sequencing submissions, GEO runs an automated, template-based validator at upload that checks required fields, controlled-list values in designated columns, and the presence of referenced files.^4^

In contrast, BioStudies supports MINSEQE through the Annotare submission system (Kolesnikov *et al*., 2015). Annotare provides structured, template-driven forms: submitters select a *technology* and a *material type*; these choices pre-select the minimum set of mandatory sample attributes. The forms offer ontology-backed suggestions for many fields and the built-in validator must be passed before curation.^5^ Submissions are also reviewed by human curators who can flag missing or inconsistent metadata. This results in more structured and semantically richer submissions compared to GEO, though gaps in coverage still exist for complex study designs.

#### Proteomics

For proteomics, the MIAPE guideline provides standardized metadata expectations for sample preparation, instrumentation, data acquisition, and analysis workflows (Taylor *et al*., 2007). One major public repository is PRoteomics IDEntifications database (PRIDE), part of the ProteomeXchange consortium, which supports MIAPE-compliant submissions through its dedicated submission tool (Perez-Riverol *et al*., 2025). The submission workflow applies semantic constraints to a subset of core metadata fields, while leaving descriptive metadata largely unconstrained. Metadata compliance is ensured through mandatory requirements, ontology-based term selection (via integrated ontology lookup services), automated validation checks within the submission pipeline, and curator review prior to dataset acceptance.

#### Metabolomics

In the case of metabolomics, the Metabolomics Standards Initiative (MSI) has proposed a set of minimum reporting standards that addresses metadata for sample handling, chemical analysis, and metabolite identification (Sumner *et al*., 2007). The MetaboLights repository serves as one of the primary submission portal for MSI-compliant metabolomics data and provides a structured, template-driven submission workflow that captures experimental design, sample metadata, and analytical methods (Yurekten *et al*., 2024). The submission system supports selected controlled vocabularies and ontology-backed fields for core metadata elements, while allowing free-text descriptions for methodological details.

While these platforms support structured submission templates and controlled vocabularies within their respective domains, their metadata models are largely developed independently and do not provide explicit semantic alignment across different omics domains or with regulatory metadata standards. Cross-domain initiatives such as the NCBI BioSample (Barrett *et al*., 2012) and EBI BioSamples databases (Courtot *et al*., 2022) provide a shared infrastructure for harmonizing biological sample descriptions across multiple omics repositories; however, this harmonization is limited to sample-level metadata and does not extend to experimental design, assay-specific metadata, or regulatory standards.

#### Regulatory Frameworks

The OORF was developed to support the regulatory acceptance of omics data by establishing modular, structured metadata requirements specific to toxicology. The framework includes modules for transcriptomics and metabolomics, while additional modules for proteomics are currently under review for adoption. OORF extends beyond generic metadata reporting by including toxicology-specific descriptors such as species selection rationale, exposure justification, and sampling schemes (OECD, 2023). However, it is not yet available in a machine-actionable format, and implementation in public repositories remains limited. Filled OORFs can be distributed as supplementary metadata files alongside datasets in existing omics repositories, including GEO, PRIDE, MetaboLights, and BioStudies. While these repositories do not interpret or validate such records, they ensure long-term preservation and contextual linkage to the data.

#### Laboratory Practices

Metadata captured in laboratory environments is often recorded in heterogeneous formats, including Electronic Lab Notebooks (ELNs), spreadsheets, lab books, and Laboratory Information Management Systems (LIMSs). At Helmholtz Centre for Environmental Research (UFZ), we assessed metadata practices across three departments (Computational Biology and Chemistry, Molecular Toxicology, and Ecotoxicology), revealing further divergence from public standards. Local documentation frequently contains toxicology-specific descriptors that are not represented in standard repository templates, such as detailed exposure justification, vehicle composition, organism acclimatization conditions, or batch-specific extraction protocols. Conversely, repository-required structured elements, such as ontology-based annotations and controlled vocabulary terms, are often absent or inconsistently applied in laboratory records. In addition, for dry-lab workflows, relevant metadata is often embedded in analysis scripts and workflow configuration files, where it remains largely implicit and is typically stored separately from wet-lab metadata.

To conclude, existing metadata frameworks provide essential foundations but lack interoperability across omics domains and are often not optimized for toxicological contexts. Most reporting frameworks are primarily designed for human readability, relying on narrative descriptions or loosely structured fields rather than fully machine-actionable representations. Our harmonization strategy builds on these standards to define a unified metadata model that aligns repository-specific formats with regulatory requirements and enriches them with domain-specific metadata tailored to institutional workflows.

### 2.2 Curating the Toxicology Metadata Pool

The first goal in the MeSyTo workflow was the construction of a metadata pool for omics-based toxicology studies. Rather than defining a metadata schema from scratch, we collected candidate metadata fields from regulatory reporting frameworks, public repository requirements, community standards, and internal laboratory workflows. The aim of this step was to capture the breadth of metadata required to describe toxicological omics experiments across different levels of granularity, including study-level information, test system characteristics, exposure conditions, individual samples, assay-specific acquisition and processing steps, and downstream data analysis metadata.

Transcriptomics was used as the primary pilot case for systematic metadata alignment. Field selection began with the available OORF modules most relevant to our use cases, including the Toxicology Experiment Reporting Modules (TERMs) to capture the overall study design, the Data Acquisition and Processing Reporting Modules (DAPRMs) for documenting RNA-Seq workflow, and the Data Analysis Reporting Modules (DARMs) for the analysis of differentially abundant molecules. These modules provided a regulatory-oriented foundation for describing study design, test systems, exposure scenarios, sequencing workflows, and analysis of differentially abundant molecules. This initial field pool was then extended with metadata elements from public transcriptomics repositories, in particular GEO and BioStudies, while excluding concepts specific to clinical or agricultural contexts that were not relevant to toxicological omics. Finally, the transcriptomics pool was complemented with metadata elements derived from internal UFZ workflows, including descriptors relevant to mechanistic toxicology classifications and ecotoxicological model systems such as zebrafish embryos, which necessitated the inclusion of fish husbandry and aquatic exposure data.

To extend the framework beyond transcriptomics, we incorporated metadata fields relevant to both metabolomics and proteomics. For metabolomics, the initial metadata set was based on the mass spectrometry–oriented OORF DAPRM module and the Metabolomics Standards Initiative. For proteomics, metadata fields were incorporated from the OORF proteomics module that is currently under development, MIAPE, and internal UFZ toxicoproteomics workflows. These omics-specific extensions ensured that the MeSyTo metadata model could represent central aspects of sample preparation, analytical acquisition, quality control, and downstream processing for mass spectrometry–based workflows.

This implementation therefore reflects an intentionally staged harmonization strategy. Transcriptomics served as the most mature alignment case and was used to develop and test the overall approach across regulatory, repository, and laboratory contexts. Metabolomics and proteomics were incorporated to reflect a multi-omics structure within the MeSyTo metadata model, but they do not yet have the same level of coverage or profile-specific implementation as transcriptomics. The resulting metadata pool thus serves both as the basis for the current harmonized schema and as an extensible foundation for future refinement as additional standards, repository requirements, and laboratory workflows are integrated.

The curated metadata pool formed the starting point for subsequent semantic harmonization (Figure 1, Step 2). In the following sections, we describe how heterogeneous metadata keys were aligned across frameworks and mapped to ontology terms, and how metadata values were addressed through controlled vocabularies, unit specifications, and normalization strategies.

### 2.3 Harmonizing and Mapping Metadata Keys Across Frameworks

Metadata field names, or keys, are frequently inconsistent across metadata frameworks. The same concept may be represented under different labels such as *organism, species*, or *animal species*, depending on the source. Conversely, similar field names may differ in their intended scope, granularity, or reporting context. Therefore, harmonization required more than lexical matching of field labels. To enable semantic interoperability, we manually curated the keys contained in the metadata pool and mapped corresponding concepts across frameworks based on their labels, definitions, descriptions, and contextual use. Where possible, harmonized concepts were linked to reference ontology terms to support interoperability and unambiguous interpretation. The EFOs and the NCIts served as primary reference sources. For domain-specific concepts not adequately covered by these resources, additional specialized ontologies were considered, provided that they offered stable identifiers, clear definitions, and sufficient community acceptance. This procedure resulted in a crosswalk between source-specific metadata fields and harmonized MeSyTo concepts. The cross-walk preserves the original terminology of each framework while enabling semantically equivalent or related fields to be interpreted through a shared representation.

In the course of this manual alignment, three recurring patterns emerged: (i) shared core fields with high semantic correspondence across frameworks, (ii) conceptually related fields with divergent granularity or structural representation, and (iii) framework- or workflow-specific fields without direct counterparts. First, shared core fields formed a common interoperable backbone of the harmonized schema including descriptors such as *organism, sex, test item, exposure duration*, and *assay*. In such cases, source-specific field names could be mapped to a common MeSyTo concept while preserving the original labels used by each framework. For example, the same biological descriptor may appear as *organism* in one repository, *species* in an internal workflow, and *animal species* in a regulatory reporting context. In several instances, however, harmonization also required structural refinement. For example, the OORF field describing the *size and type of sequencing* (e.g., 100 bp paired-end) combines information on read length and library layout. To align this descriptor with the BioStudies *library layout* field and the GEO *single or paired-end* field, the composite attribute was decomposed into two subfields: *Size of sequencing* and *Type of sequencing*, thereby preserving OORF terminology.

Second, many concepts were related across frameworks but differed in granularity or structural representation. In these cases, harmonization required more than assigning equivalent labels. For example, in exposure experiments, GEO largely represents treatment information through free-text *treatment protocol* fields and sample-level *treatment* descriptions. BioStudies similarly combines protocol-level descriptions with more structured sample attributes such as *compound, dose*, and *exposure time*. In contrast, OORF decomposes exposure information into a more detailed set of structured elements, reflecting its emphasis on regulatory traceability. While the entire OORF exposure section could technically be mapped to a single higher-level concept such as *treatment protocol*, such a many-to-one consolidation would collapse critical semantic distinctions and undermine interoperability at this stage of harmonization. Moreover, OORF captures exposure metadata primarily as summary information linked to the overall study design, whereas GEO and BioStudies collect much of this information at the level of individual samples. Although sample-level metadata can be submitted to OORF as a separate “data object,” the harmonized schema needed to preserve these structural differences by distinguishing study-level summaries, exposure-level descriptors, and sample-level annotations.

Third, several metadata fields were specifically tailored for individual frameworks or use cases. The OORF, for example, includes regulatory descriptors such as *test guideline compliance, study rationale*, and provenance information that are not typically represented in general-purpose repositories. Conversely, repository-specific submission fields may be required for data deposition but do not necessarily correspond to regulatory reporting elements. Internal UFZ workflows added further descriptors reflecting local experimental practice, including model-specific *husbandry* information or laboratory-specific processing details. These framework- or workflow-specific fields were retained when they contributed to toxicological interpretation, regulatory transparency, repository submission, or cross-laboratory reuse.

Together, these three patterns illustrate that harmonization cannot rely on naïve one-to-one field equivalence but instead requires a combination of direct semantic alignment, structural decomposition, concept-level consolidation, and retention of context-specific metadata fields. Therefore, the harmonized MeSyTo concepts were not treated as flat field aliases, but represented as ontology-backed properties within a structured metadata model, enabling both source-specific terminology and shared semantic interpretation.

### 2.4 Harmonizing Metadata Values

Compared to the limited set of metadata keys, metadata values pose a greater harmonization challenge: they are numerous, heterogeneous, and frequently expressed in free text. Across datasets, the same entity may be described using different terms, abbreviations, or formats. For example, the species *Mus musculus* may be denoted as *house mouse* or described more specifically as *mouse (C57BL/6)*, whereas an exposure duration of *1 day* may also be recorded as *24 h* or *24 hours*. Such variation prevents automated comparison and reuse of datasets and necessitates labor-intensive manual curation. Linking values to controlled vocabularies, ontology concepts, and standardized unit representations enables synonym resolution, reduces ambiguity, and supports querying and integration across studies. Thus, value-level harmonization is a critical step toward machine-actionable interoperability.

The treatment of metadata values varies significantly between platforms. The OORF adopts a largely free-text approach, offering maximum flexibility but relying on user expertise for consistent terminology, formatting, and level of detail. GEO follows a hybrid strategy: spreadsheet-based submissions include controlled lists for selected technical columns, such as *molecule, library strategy*, and *instrument model*, and organism names are validated against NCBI Taxonomy, whereas most descriptive fields remain free text. EBI BioStudies, via Annotare, implements a more semantically guided model, using dropdown menus for technical attributes and ontology-backed suggestions, primarily from EFO and related ontologies, for biological descriptors such as *organism, sex*, and *disease*. It also supports unit specification for numerical fields and allows custom attributes with optional ontology linking. Other omics repositories show similar combinations of structured submission, controlled terminology, and remaining free-text flexibility. In proteomics, PRIDE submissions are guided through the PRIDE/PX Submission Tool and include required dataset metadata, controlled vocabulary support, and ontology lookup for selected annotations. In metabolomics, MetaboLights uses a structured submission workflow based on study, sample, and assay metadata, with validation prior to submission. Thus, existing platforms already provide partial mechanisms for value standardization, but they differ in which fields are constrained, which vocabularies are used, and how much descriptive metadata remains unconstrained.

In the course of this value-level assessment, three recurring patterns emerged: (i) fields for which existing framework-specific standards can be enforced directly, (ii) complex biological or toxicological values that require ontology-based mapping with pragmatic curation, and (iii) descriptive or identifier fields for which guidance is more appropriate than strict restriction. First, some metadata fields are already associated with controlled value sets defined by the target framework or repository. For these fields, value harmonization can rely on enforcing the respective existing standard. This applies, for example, to technical descriptors such as *library source*, as well as to simple categorical descriptors such as *sex*, where values can be restricted to an authoritative vocabulary to ensure consistency. However, even for apparently similar fields, permissible values may differ between platforms. For example, because repositories broker sequencing submissions to different archives, GEO to Sequence Read Archive (SRA) and BioStudies to European Nucleotide Archive (ENA), their accepted controlled vocabularies may diverge.

Second, many biological and toxicological values require ontology-based mapping combined with pragmatic manual curation. Concepts such as *tissue, chemical exposure*, or *organism* may appear as free-text labels, synonyms, abbreviations, or at different levels of specificity across datasets. Mapping these values to resources such as UBERON, ChEBI, or NCBI Taxonomy supports semantic precision and cross-study querying, while practical curation rules are needed to handle synonyms, strain information, term aggregation, and unit representation.

Third, some metadata values cannot be usefully restricted to closed vocabularies. This is particularly true for descriptive fields, protocol summaries, comments, and certain identifier fields, where excessive restriction would reduce expressiveness or prevent the reporting of relevant context. For such fields, value harmonization should rely on guidance rather than enforcement. Recommended templates, formatting conventions, examples, and optional ontology tagging can promote consistency while preserving the flexibility required for complex toxicological experiments.

Together, these patterns show that value-level harmonization requires a combination of controlled vocabulary enforcement, ontology-based normalization, and flexible guidance for free-text fields. Figure 2 provides a schematic illustration of metadata harmonization, using organism annotation as an example to show how heterogeneous framework-specific representations can be mapped to a shared semantic reference.

**Figure 2:**
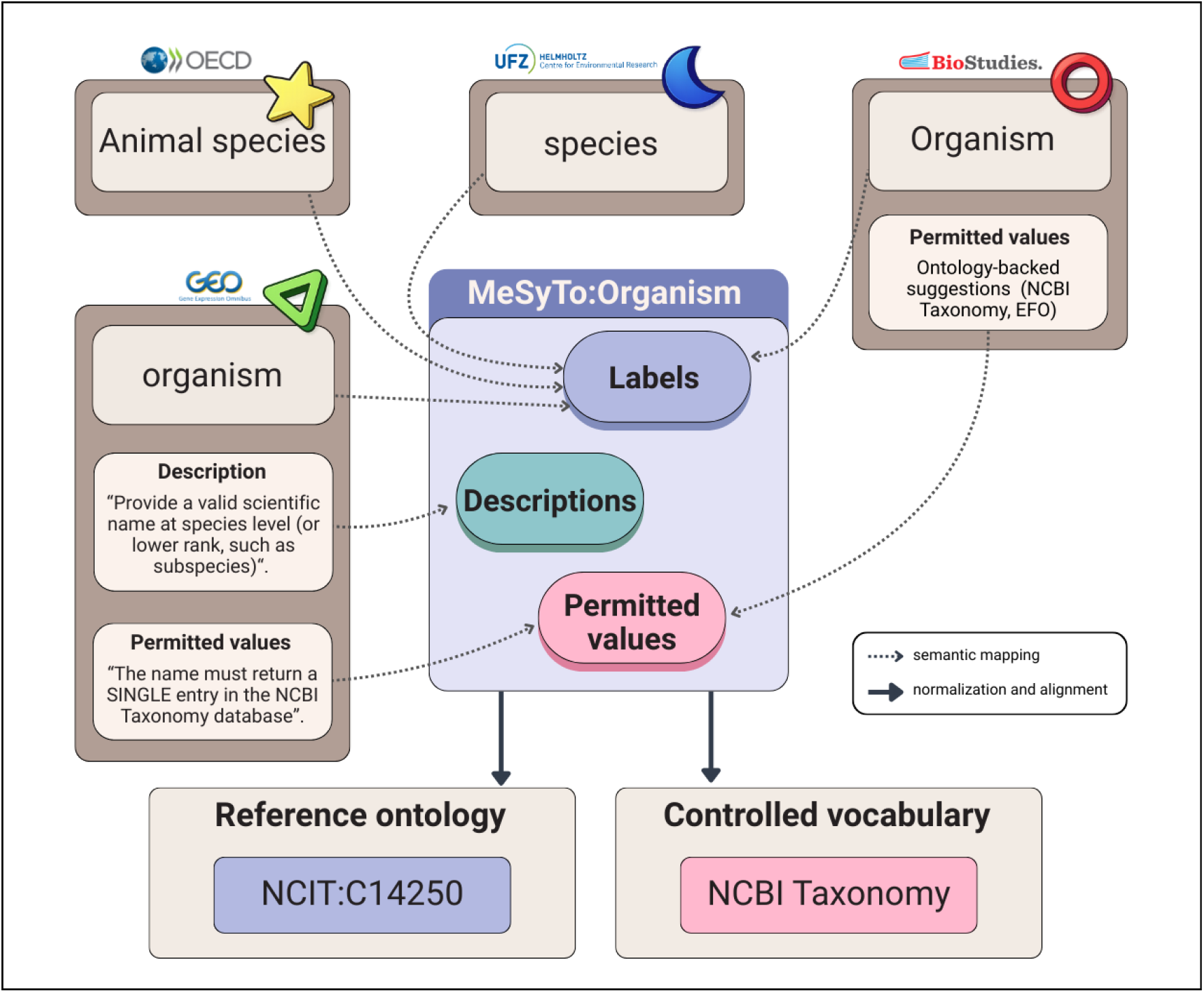
Metadata harmonization across frameworks. Example of key- and value-level harmonization using the *Organism* concept. Semantically equivalent metadata fields from different sources are mapped to a shared MeSyTo concept while preserving source-specific labels, descriptions, and permitted values. Where possible, metadata values are normalized through controlled vocabularies and aligned to reference ontology terms to support consistent interpretation across frameworks.

## 3 Metadata Model Design and Validation

To operationalize the harmonization of toxicological omics metadata described above, we implemented MeSyTo as a semantic metadata model and validation framework for machine-actionable metadata. While machine-readable metadata can be parsed by software, machine-actionable metadata additionally requires explicit semantics and constraints that enable validation, transformation, and reuse across repository, regulatory, and institutional contexts. In the following sections, we describe the design of the MeSyTo Metadata Model (MMM) and its implementation through SHACL-based validation profiles and metadata collection forms.

### 3.1 MeSyTo Metadata Model Development

As depicted in Step 3 of Figure 1, we developed the MMM in `Protégé` (Musen, 2015), an open-source ontology editor and knowledge management system widely used in the Semantic Web community. The aim of the MMM was to provide a semantically consistent representation of metadata fields relevant to toxicological omics and to implement the harmonized concepts and mappings in Section 2 in a machine-readable form. `Protégé` provides a graphical environment for creating and managing knowledge models represented in RDF, the W3C standard for machine-readable data^6^. RDF represents knowledge as statements, or *triples*, that link resources through defined relationships. In this framework, entities such as a *sample*, a *treatment compound*, or the relationship between them can be identified by unique Internationalized Resource Identifiers (IRIs). These identifiers support standardization by avoiding naming conflicts and enable provenance tracking by allowing the origin of metadata statements to be referenced unambiguously.

The MMM is formalized in OWL, which builds upon RDF and RDF Schema (RDFS). While RDF provides the basic triple-based structure and RDFS introduces vocabulary for defining classes and properties, OWL allows richer semantic relationships to be represented. In the MMM, OWL is used primarily to define the conceptual structure of the metadata model, including classes, properties, labels, and links to external ontology terms. Constraint enforcement and profile-specific validation are implemented separately through SHACL, as described in Section 3.2. All MMM elements are defined as RDF resources under the mesyto: namespace (https://w3id.org/mesyto/ontology#), ensuring globally consistent identifiers across tools and exports. The MMM implementation is publicly available^7^.

The MMM serves as a centralized knowledge model that captures the metadata fields relevant to omics-based toxicology. Its class hierarchy is conceptually aligned with the modular logic of the OORF: top-level classes reflect major reporting domains inspired by OORF modules, while subordinate classes organize these domains into thematic sections and subsections. Collectively, these classes cover three principal domains: (i) toxicology experiment—related descriptors, including test item descriptors, test system descriptors, and treatment conditions; (ii) omics technology— related descriptors and upstream analysis parameters covering *transcriptomics, proteomics*, and *metabolomics*; and (iii) data analysis-–related descriptors, including the software, parameters, inputs, and outputs relevant for data interpretation and reproducibility. Figure 3(A) depicts the high-level organization of metadata classes.

**Figure 3:**
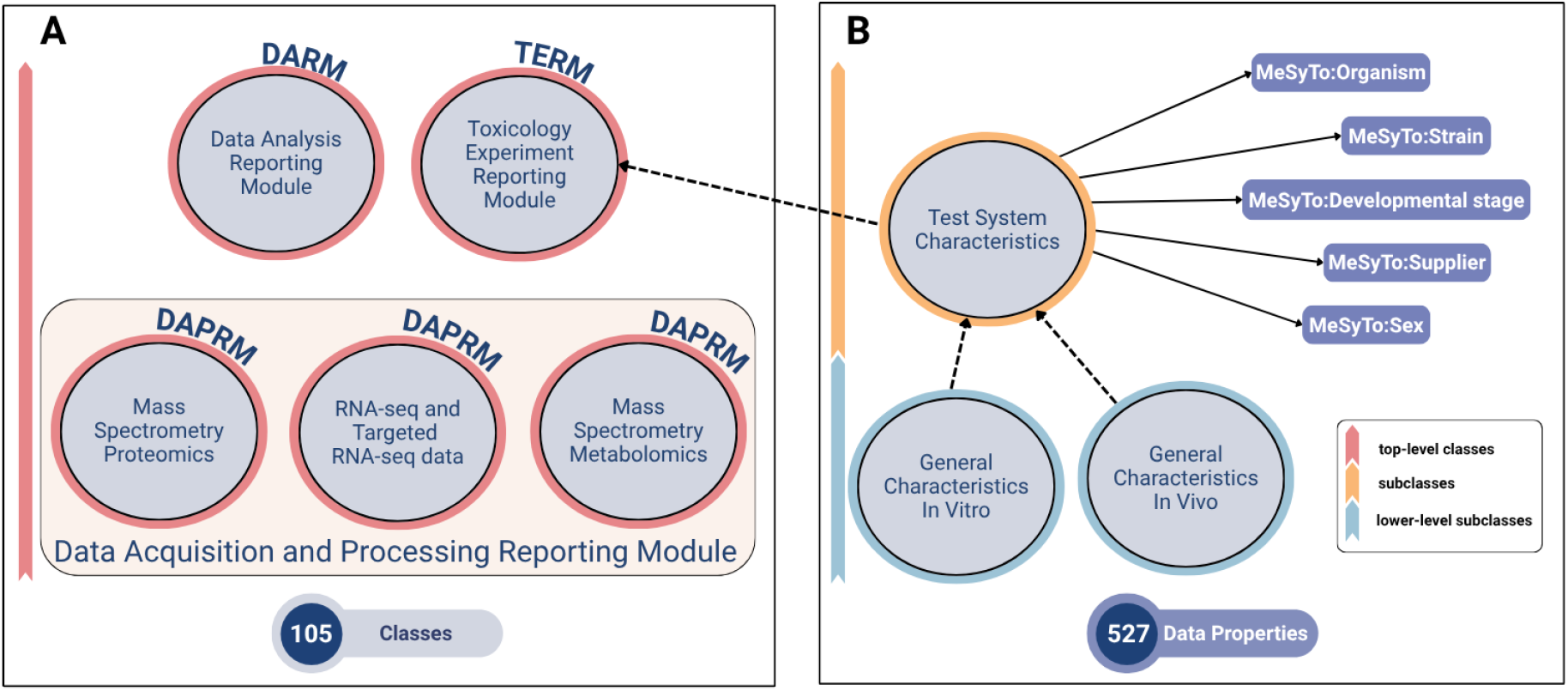
Overview of the MeSyTo Metadata Model. (A) Top-level organization of the MMM into toxicological experiment-, omics technology-, and data analysis-related domains, following the modular structure of the OORF. The model spans multiple omics layers, including metadata fields for RNA-Seq and mass spectrometry–based techniques as proteomics and metabolomics. (B) Example MMM class illustrating test system characteristics and associated data properties corresponding to harmonized MeSyTo concepts. Source-specific labels, descriptions, and ontology mappings are retained at the property level, while datatype constraints and permitted values are implemented in SHACL shapes rather than enforced directly in the MMM. In total, the current implementation contains 105 classes and 527 data properties. For clarity, only selected classes, subclasses, and data property relationships are shown.

Within these classes, harmonized MeSyTo concepts are represented as properties, specifically data properties in OWL. These properties implement the crosswalk between source-specific metadata fields and shared MeSyTo concepts. They are annotated with alternative labels, definitions, framework-specific requirement levels, external ontology mappings, controlled vocabularies, and constraint-relevant information, such as expected datatypes or cardinality. In this way, the MMM preserves source-specific terminology while providing a shared semantic layer for consistent metadata annotation. In addition, properties can be assigned MeSyTo-specific requirement levels and vocabulary constraints, which are subsequently translated into SHACL validation rules for metadata considered essential for toxicological interpretation and reuse, even when such information is optional or unconstrained in the target repositories.

As a metadata schema, the MMM does not store experimental data itself but defines the structure and semantics required for its annotation. By separating toxicological experiment descriptors from omics assay technologies and bioinformatics analysis components, the MMM enables the reuse of entities across multi-omics studies. For example, a single toxicological experiment may be associated with multiple omics assays, while comparable bioinformatics analyses can be applied to different omics layers, sharing common descriptors and differing only in technology-specific parameters.

An example MMM class illustrating test system characteristics and the corresponding data properties used for metadata recording is shown in Figure 3(B). The harmonized concept *MeSyTo:Organism*, previously introduced in Figure 2, is represented here as a data property of the test system characteristics class. This illustrates how a source-specific metadata field is represented within the common semantic structure of the MMM, while still retaining its original labels and descriptions as well as mappings to external ontology terms.

In the context of interdisciplinary toxicology projects, the MMM helps to resolve terminological ambiguities and supports consistent metadata annotation across diverse experimental settings. In total, it currently comprises 105 classes and 527 data properties, yet remains extensible to incorporate new terms, additional metadata sources, and emerging omics technologies. Versioned releases of the MMM are archived on Zenodo.^8^

### 3.2 SHACL-based Validation and Data Collection

To support guided acquisition of toxicological omics metadata and to ensure completeness and consistency, the MeSyTo workflow employs SHACL ^9^ in combination with SKOS, both W3C standards for working with RDF-based data (Figure 1, Step 4). SHACL enables the formal definition of validation shapes over the classes and properties defined in the MMM. These shapes specify which metadata fields are expected or required and how their values should be represented, including datatypes, cardinalities, controlled vocabularies, string patterns, and other validation constraints. SKOS is used to represent controlled vocabularies in a reusable and machine-readable form, allowing permitted values to be maintained separately from the structural metadata model while remaining linked to the corresponding SHACL constraints. This validation principle is illustrated schematically in Figure 4, where profile-specific SHACL shapes are shown as filters that constrain metadata according to the requirements of different target frameworks.

**Figure 4:**
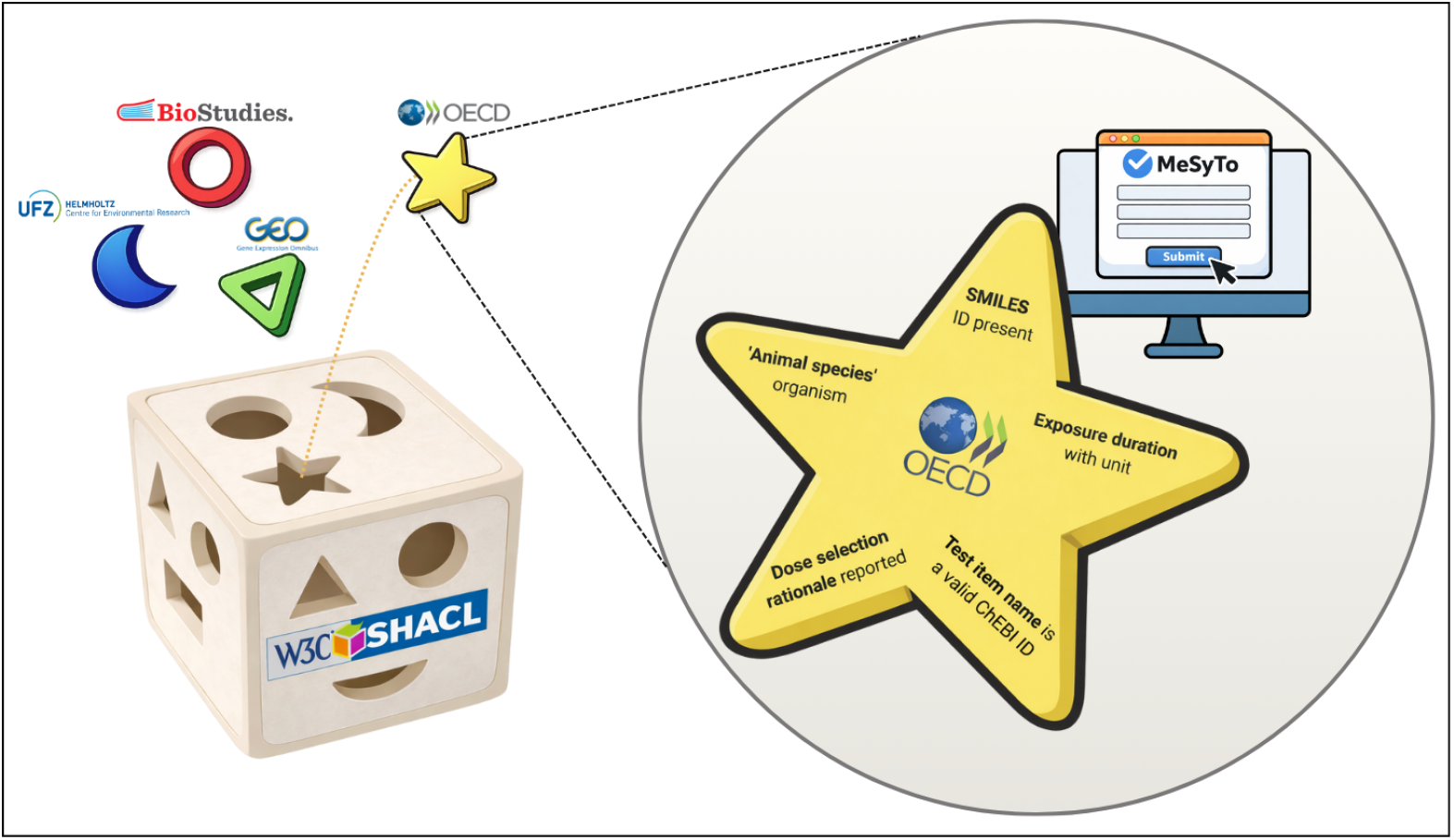
SHACL-based profile validation and form generation. Schematic representation of the SHACL validation layer used to derive and enforce profile-specific metadata requirements. Each opening represents a target profile, such as GEO or BioStudies, through which only metadata conforming to the corresponding SHACL constraints can pass. The magnified OORF example illustrates how profile-specific labels, required fields, controlled vocabularies, datatype constraints, and unit requirements are encoded. The same profile shapes can also be used to generate metadata collection forms.

The model-to-forms pipeline proceeds in three steps. First, reusable base shapes and controlled vocabularies are derived from the MMM. Second, these base shapes are transformed into profile-specific SHACL shapes for individual target frameworks, such as GEO, BioStudies, OORF, or internal UFZ metadata exchange. Third, the profile shapes are supplemented with form-specific information, such as field order, section headings, grouping, and display labels, and are rendered as interactive forms for metadata collection.

#### Base shapes and vocabularies

To manage the large and heterogeneous set of metadata fields collected across the MeSyTo project and to provide a reusable validation layer, we first generated base SHACL shapes from the OWL implementation of the MMM using a dedicated script ^10^. These base shapes mirror the top-level classes of the MMM, including TERM, DAPRM for RNA-seq, proteomics, and metabolomics, and DARM, resulting in five base shapes. They contain the complete set of metadata fields defined in the MMM together with the framework-specific annotations encoded for each property. Rather than being used directly as user-facing forms, the base shapes provide a common foundation for constructing profile-specific shapes.

During base shape generation, information represented in the MMM is translated into executable SHACL constraints. For example, property ranges are converted into datatype constraints, such as string, integer, decimal, boolean, or date, while cardinality and requirement information are translated into corresponding validation rules. In parallel, permitted values identified during metadata curation are converted into SKOS vocabularies,^11^ which are generated using a dedicated script.^12^ These vocabularies are maintained independently of the structural model and can be referenced by SHACL constraints wherever controlled value selection is required.

#### Profile shapes

As the second step of the pipeline, profile-specific SHACL shapes are generated from the base shapes using a dedicated script^13^. Whereas base shapes contain the complete set of MMM-derived metadata fields, profile shapes represent framework- or use-case-specific subsets tailored to a particular target context. In the current MeSyTo implementation, we provide four profiles: metadata preparation for submission to GEO and BioStudies, metadata required for OORF regulatory reporting, and metadata for internal data exchange within UFZ (Figure 1, Step 5).

Profile shapes are derived by selecting the fields relevant to the corresponding target framework or use case. During this process, labels and descriptions are adapted to the terminology of the target framework, while annotations irrelevant to that framework are omitted. Framework-specific and MeSyTo-specific requirement levels, controlled vocabulary bindings, and additional validation constraints are translated into SHACL. Thus, each profile preserves the shared semantic structure of the MMM while enforcing the metadata requirements and terminology of a specific reporting or submission context.

When multiple frameworks provide different controlled vocabularies for the same metadata concept, each profile can retain its native vocabulary. Profiles that would otherwise rely on free-text values can instead reuse a selected canonical vocabulary defined in the MMM. Canonical vocabularies are inherited by profile-specific shapes unless a profile explicitly defines an alternative vocabulary, which then takes precedence.

#### Forms for metadata collection

As the final step of the pipeline, profile-specific shapes are transformed into user-facing metadata collection forms. To support this transformation, the profile shapes are supplemented with form-specific information, including field order, section headings, grouping, display labels, and explanatory text. This allows the same underlying MeSyTo concepts to be presented differently depending on the target profile, reflecting the reporting logic and terminology of frameworks such as GEO, BioStudies, OORF, or internal UFZ workflows.

As a proof of concept, we implemented a prototype web application using the *shacl-form* HTML5 web component (ULB Darmstadt, 2023), which renders interactive forms from SHACL shapes and serializes user input as RDF compliant with the underlying metadata model. The application demonstrates how MeSyTo profile shapes can be used to generate framework-specific metadata collection forms while preserving a shared semantic representation in the background. A demo version of the application is available from the public MeSyTo GitLab repository.^14^ The application supports both native SHACL-based validation and additional JavaScript-enhanced checks. Native SHACL constraints include datatype validation, cardinality constraints, required fields, controlled vocabulary restrictions, string-length restrictions, and pattern matching. JavaScript extensions are used for interactive functionality that is difficult to express in static SHACL shapes, such as dynamic field dependencies, custom conditional checks, interactive user feedback, and lookups against external semantic resources such as ChEBI and NCBI Taxonomy.

## 4 Scope, Limitations, and Future Development

### Scope

The current implementation of MeSyTo provides a functional foundation for ontology-driven metadata harmonization in toxicological omics from start-to-finish. Its most mature application is in the context of transcriptomics, where the framework supports metadata collection and validation for repository submission preparation, regulatory reporting, and cross-laboratory data exchange. This reflects the staged development strategy described above: transcriptomics was used as the primary alignment case because it provided the strongest overlap between established repository formats, OORF reporting requirements, and internal UFZ workflows. At the same time, the MMM already incorporates metadata elements relevant to mass-spectrometry-related high-throughput techniques such as proteomics and metabolomics, thereby establishing a multi-omics structure that can be extended as additional standards, repository requirements, and laboratory workflows are integrated. In its present form, MeSyTo therefore does not replace existing repositories or reporting frameworks, but provides a semantic coordination layer that links their terminology, preserves source-specific requirements, and supports the generation of context-specific validation profiles. This represents a substantial harmonization effort, as the MMM maps metadata concepts across frameworks with different purposes, terminologies, and levels of granularity, ranging from study- and experiment-level descriptions to exposure, assay, and sample-level annotations. The presented prototype includes the MMM, SKOS vocabularies, SHACL base shapes, profile-specific shapes for selected target contexts, and a proof-of-concept form-based metadata collection workflow. All components are publicly available as open-source resources, enabling reuse, inspection, adaptation, and extension by the toxicological omics community. The ontology, validation shapes, controlled vocabularies, and associated scripts are maintained in the public MeSyTo GitLab repository under version control, with versioned releases archived on Zenodo. Community feedback, feature requests, and issue reporting are supported through the GitLab issue tracker.

### Limitations

A central limitation of the current ecosystem is that public omics repositories are not (yet) designed to fully capture, validate, or expose the breadth of metadata required for toxicological multi-omics studies. Repositories such as GEO, BioStudies, and PRIDE support broad omics data deposition and allow submitters to provide custom sample attributes; however, these fields remain only partly constrained and are not standardized. Depending on the repository, sample attributes may additionally be annotated with units and ontology terms, whereas others rely more strongly on free-text representations. Nevertheless, these mechanisms mainly address sample-level metadata and do not comprehensively cover study-level regulatory descriptors, detailed exposure metadata, assay-specific processing information, or dry-lab workflow and analysis metadata. MeSyTo can prepare harmonized and semantically annotated metadata for such contexts, but its full value will depend partly on whether repositories or downstream platforms can ingest, preserve, and expose harmonized labels, controlled vocabularies, ontology mappings, and other semantic annotations. As a full standardization of toxicological omics metadata remains a long-term goal, MeSyTo currently functions primarily as a complementary harmonization and validation layer rather than as a replacement for repository submission systems. Finally, SHACL-based validation can check structural and formal consistency, such as required fields, datatypes, value restrictions, and controlled vocabulary use, but it cannot by itself guarantee the scientific correctness of the submitted metadata.

### Future perspectives

Future development of MeSyTo will focus on extending the framework from a prototype for harmonized metadata collection and validation toward a reusable infrastructure for toxicological omics metadata management. One important direction is the closer alignment of MeSyTo profiles with public repository submission workflows. Although direct adoption of MeSyTo by repositories would provide the strongest route toward standardized toxicological omics metadata, the framework can also provide practical support without requiring changes to repository infrastructures. For example, MeSyTo profiles could be used to generate repository-specific metadata templates, pre-filled submission spreadsheets, or guidance documents that help researchers prepare metadata for resources such as GEO, BioStudies, PRIDE, and MetaboLights. Where supported by repository interfaces, these exports could be further extended toward semi-automated submission workflows. In parallel, MeSyTo could serve as a complementary metadata layer that preserves ontology mappings, harmonized labels, controlled vocabularies, and validation results even when the target repository cannot represent these annotations directly.

A second development direction is the integration of MeSyTo with institutional metadata infrastructures, in particular LIMSs, ELNs, workflow management systems, and institutional research data management platforms. In practice, such systems are often the primary sources of sample tracking information, experimental protocols, instrument records, workflow configuration, and laboratory-specific metadata. Closer integration would allow existing sample- and experiment-level information to be reused during metadata preparation, reducing redundant manual entry and improving consistency between laboratory documentation, repository submissions, and regulatory reporting. This would be particularly valuable for toxicological omics workflows, where metadata describing test systems, exposure conditions, sampling schemes, analytical processing, and computational analysis is distributed across wet-lab and dry-lab environments. Recent developments in ELN and research data management interoperability illustrate how such integration could be implemented. For example, `ELNdataBridge` (Starman *et al*., 2025), provides a server-based adapter architecture for exchanging and synchronizing data between different ELN platforms through Python APIs. Complementary approaches based on standardized exchange formats, such as the Research Object Crate (RO-Crate)-based `.eln` format (Soiland-Reyes *et al*., 2022), could support transfer of semantically annotated experiment records, including references to external ontology terms. API-enabled platforms such as `eLabFTW` (Hewera *et al*., 2021) or `openBIS` (Barillari *et al*., 2016) further show how laboratory documentation, inventory management, and research data management systems can be accessed programmatically and integrated with external services. Building on these patterns, a MeSyTo-based API could allow metadata records to be submitted/collected for validation, transformed between profiles, enriched with ontology mappings, and exported in repository-specific or RDF-based formats. Such an interface would support automated exchange between MeSyTo, institutional ELN/LIMS environments, data portals, and workflow systems, allowing the MeSyTo validation layer to operate as part of existing research data management workflows rather than as a separate step.

Large Language Models (LLMs) offer promising opportunities to further extend the MeSyTo framework. In particular, LLM-based approaches could help bridge differences in reporting granularity between metadata frameworks by enabling bidirectional translation between structured metadata fields and free-text protocol descriptions. This would facilitate seamless transitions between profiles that rely on narrative reporting and those that require highly structured parameterization. In addition, LLMs could support semi-automated metadata acquisition by extracting relevant information from workflow configuration files, manuscript drafts, and ELNs. When combined with SHACL-based validation, such approaches could substantially reduce manual effort while preserving semantic consistency, traceability, and validation guarantees. However, such approaches would require human-in-the-loop review, with SHACL validation serving as a formal quality-control layer rather than as a replacement for expert curation.

In the longer term, MeSyTo could provide the basis for a standalone metadata registry or database for toxicological omics studies. This would help preserve semantic annotations that are currently difficult to represent in general-purpose repositories and would support cross-study querying, reuse, and comparison across omics layers, experimental systems, and regulatory contexts. Such an infrastructure would not replace primary data repositories, but would complement them by storing harmonized, ontology-linked metadata records and persistent references to the corresponding deposited datasets.

Together, these developments would strengthen MeSyTo as an extensible, community-oriented framework for making toxicological omics metadata more consistent, interoperable, and machine-actionable.

## RNA-seq Metadata Case Study

To demonstrate the practical utility of MeSyTo, we applied the MMM to RNA-seq metadata from NCBI BioProject PRJNA695243, a thyroid toxicity study reported by Canzler et al. (Canzler *et al*., 2025). GEO sequencing submission fields were mapped to MMM data properties and compared with structured fields retrieved from NCBI BioSample, SRA, and RunInfo. The complete methods, mapping table, and results are provided in the Supplementary Information archived on Zenodo.^15^

## Acknowledgements

The MeSyTo project was funded 2024–2026 by the Initiative and Networking Fund of the Helmholtz Association (Helmholtz Metadata Collaboration (HMC), Project Cohort 2023). We thank Alexandra Schaffert (FHAIVE – Finnish Hub for Development and Validation of Integrated Approaches, Tampere University, Finland) and Ksenia Groh (Eawag, Swiss Federal Institute of Aquatic Science and Technology, Switzerland) for sharing draft versions of the OORF proteomics reporting modules currently under review at the OECD level.

## Footnotes

1 https://www.ncbi.nlm.nih.gov/geo/

2 https://www.ebi.ac.uk/biostudies/

3 https://www.ebi.ac.uk/pride/

4 https://www.ncbi.nlm.nih.gov/geo/info/validation.html

5 https://www.ebi.ac.uk/fg/annotare/help/getting_started.html; https://www.ebi.ac.uk/fg/annotare/help/sample_attributes.html; https://ebi-gene-expression-group.github.io/annotare-help/pages/validate_exp.html

6 https://www.w3.org/standards/techs/rdf

7 https://zenodo.org/records/21033443/files/mesyto.ttl

8 https://doi.org/10.5281/zenodo.21033443

9 https://www.w3.org/TR/shacl/

10 https://zenodo.org/records/21033443/files/base_shapes.zip

11 https://zenodo.org/records/21033443/files/controlled_vocabularies.zip

12 https://zenodo.org/records/21033443/files/generate_vocab.py

13 https://zenodo.org/records/21033443/files/generate_profile_shapes.py

14 https://codebase.helmholtz.cloud/department-computational-biology/software/mesyto

15 https://zenodo.org/records/21931025/files/MeSyTo_supplement.pdf

## References

Ankley, G.T. et al. (2010). Adverse outcome pathways: a conceptual framework to support ecotoxicology research and risk assessment. Environ Toxicol Chem, 29(3), 730–41.

Barillari, C. et al. (2016). openbis eln-lims: an open-source database for academic laboratories. Bioinformatics, 32(4), 638–40.

Barrett, T. et al. (2012). BioProject and BioSample databases at NCBI: Facilitating capture and organization of metadata. Nucleic Acids Research, 40(D1), D57–D63.

Brazma, A. et al. (2012). MINSEQE: Minimum Information about a high-throughput Nucleotide SeQuencing Experiment - a proposal for standards in functional genomic data reporting.

Brockmeier, E.K. et al. (2017). The role of omics in the application of adverse outcome pathways for chemical risk assessment. Toxicol Sci, 158(2), 252–262.

Buesen, R. et al. (2017). Applying ‘omics technologies in chemicals risk assessment: Report of an ecetoc workshop. Regul Toxicol Pharmacol, 91 Suppl 1(Suppl 1), S3–S13.

Canzler, S. et al. (2020). Prospects and challenges of multi-omics data integration in toxicology. Archives of Toxicology, 94(2), 371–388.

Canzler, S. et al. (2025). Evaluating the performance of multi-omics integration: a thyroid toxicity case study. Arch Toxicol, 99(1), 309–332.

Courtot, M. et al. (2022). BioSamples database: FAIRer samples metadata to accelerate research data management. Nucleic Acids Research, 50(D1), D1500–D1507.

Harrill, J.A. et al. (2021). Progress towards an OECD reporting framework for transcriptomics and metabolomics in regulatory toxicology. Regulatory Toxicology and Pharmacology, 125, 105020.

Hewera, M. et al. (2021). elabftw as an open science tool to improve the quality and translation of preclinical research. F1000Res, 10, 292.

Joseph, P. (2017). Transcriptomics in toxicology. Food Chem Toxicol, 109(Pt 1), 650–662.

Kolesnikov, N. et al. (2015). ArrayExpress update—simplifying data submissions. Nucleic Acids Research, 43(Database issue), D1113–D1116.

Madeira, C. and Costa, P.M. (2021). Proteomics in systems toxicology. Adv Protein Chem Struct Biol, 127, 55–91.

Musen, M.A. (2015). The protégé project: A look back and a look forward. AI Matters, 1(4), 4–12.

Nijsse, B., Schaap, P.J. and Koehorst, J.J. (2023). FAIR data station for lightweight metadata management and validation of omics studies. GigaScience, 12, giad014.

OECD (2023). OECD Omics Reporting Framework (OORF): Guidance on Reporting Elements for the Regulatory Use of Omics Data from Laboratory-Based Toxicology Studies. OECD Series on Testing and Assessment. OECD.

Olesti, E. et al. (2021). Approaches in metabolomics for regulatory toxicology applications. Analyst, 146(6), 1820–1834.

Perez-Riverol, Y. et al. (2025). The pride database at 20 years: 2025 update. Nucleic Acids Res, 53(D1), D543–D553.

Reardon, A. et al. (2023). From vision toward best practices: Evaluating. Front Toxicol, 5, 1194895.

Sauer, U.G. et al. (2017). The challenge of the application of ‘omics technologies in chemicals risk assessment: Background and outlook. Regulatory Toxicology and Pharmacology, 91, S14–S26.

Soiland-Reyes, S. et al. (2022). Packaging research artefacts with ro-crate. Data Science, 5(2), 97–138.

Starman, M. et al. (2025). Elndatabridge: facilitating data exchange and collaboration by linking electronic lab notebooks via api. J Cheminform, 17(1), 86.

Sumner, L.W. et al. (2007). Proposed minimum reporting standards for chemical analysis Chemical Analysis Working Group (CAWG) Metabolomics Standards Initiative (MSI). Metabolomics: Official Journal of the Metabolomic Society, 3(3), 211–221.

Taylor, C.F. et al. (2007). The minimum information about a proteomics experiment (MIAPE). Nature biotechnology, 25(8), 887–893.

ULB Darmstadt (2023). Shacl-form: HTML5 web component for editing and viewing RDF data conforming to SHACL shapes.

Wilkinson, M.D. et al. (2016). The FAIR Guiding Principles for scientific data management and stewardship. Scientific Data, 3(1), 160018.

Yurekten, O. et al. (2024). Metabolights: open data repository for metabolomics. Nucleic Acids Res, 52(D1), D640–D646.

