## Supplementary information for "From Lab Notes to Linked Data: MeSyTo for Ontology-Driven Metadata in Toxicological Omics" (from another version of this preprint: version 1, 2026-08-18)

### 1 Practical utility of the MeSyTo Metadata Model for RNA sequencing metadata

To demonstrate the practical utility of the MeSyTo Metadata Model (MMM), we applied it to RNA-seq metadata deposited under NCBI BioProject accession PRJNA695243[1], from a study investigating thyroid toxicity (Canzler *et al.*, 2025).

#### 1.1 Input data collection and preparation

We used the GEO sequencing submission spreadsheet[2] to obtain the GEO field names. These fields were matched to MMM data properties using the GEO labels included in the model.

We downloaded metadata for NCBI BioProject PRJNA695243[1] from BioSample and SRA. The metadata were retrieved as XML files using the NCBI Entrez Direct command-line tools[3]. Run-level metadata were downloaded as a CSV file from the SRA Run Selector[4]. The downloaded records were then filtered to include only RNA-seq data.

The input data consisted of three files: a BioSample XML file containing sample information, an SRA XML file containing study and experiment information, and a RunInfo CSV file containing run-level information. The filtered data contained 150 linked BioSample records, 150 SRA experiment records, and 150 RunInfo records. The field inventory contained 225 source-specific entries: 39 from BioSample, 147 from SRA, and 39 from RunInfo. Some information occurred in more than one source.

[1] https://www.ncbi.nlm.nih.gov/bioproject/PRJNA695243
[2] https://www.ncbi.nlm.nih.gov/geo/info/seq.html#metadata
[3] https://www.ncbi.nlm.nih.gov/books/NBK179288/
[4] https://www.ncbi.nlm.nih.gov/Traces/study/?acc=PRJNA695243

We then created a mapping table linking GEO fields, MMM data properties, and fields found in the downloaded NCBI metadata. A GEO field could map to more than one MMM data property when the MMM described the information in greater detail. Fields used only for file submission were excluded from the mapping table.

The complete reproducible workflow—including the downloaded input metadata, filtered intermediate files, mapping rules, scripts, and generated outputs—is available in the MeSyTo GitLab repository under the `v1.1.0` release tag.[5]

### 1.2 Mapping results

The mapping table covered 32 distinct section-specific GEO fields. Of these, 28 fields (87.5%) mapped to the MMM, while four had no MMM-specific mapping. Twenty-seven GEO fields mapped to one MMM data property each. The remaining GEO field, *data processing step*, mapped to five more specific MMM data properties describing different processing and filtering operations. The table therefore contained 36 rows: 32 GEO–MMM mappings and four unmapped GEO fields.

Beyond field-level coverage, the MMM preserves the constraints specified by GEO for the 28 mapped fields, including requirement levels and controlled-vocabulary specifications. It also provides more explicit machine-actionable constraints for datatypes, cardinalities, and permitted values. Where GEO leaves values unconstrained, the MMM can introduce an appropriate controlled vocabulary to improve consistency. For example, the MMM defines permitted values for the *sex* field rather than leaving it as unconstrained free text. In addition, fields that are optional in GEO but important for toxicological omics studies, such as *dose* and *time*, can be defined as mandatory in an MMM validation profile.

Four GEO fields lacked an MMM-specific mapping. The generic fields *contributor* and *description* can be represented using established Dublin Core terms. The other two fields are domain-specific: *ChIP antibody* is not applicable to RNA-seq, while *tumor stage* is primarily relevant to oncology studies and lies outside the current scope of the MMM.

Of the 28 GEO fields covered by the MMM, 16 (57.1%) had corresponding structured fields in the retrieved NCBI metadata for PRJNA695243. The remaining 12 fields were covered by the MMM but were not found as structured NCBI metadata. Fields concerning *cell line*, *cell type*, *disease state*, *genetic modification*, and *genotype* were not relevant to this *in vivo* study in male Wistar rats. Information on the experimental design, laboratory and data-processing protocols, and related technical details was provided in the associated publication (Canzler *et al.*, 2025), but was not available in structured form in the retrieved NCBI metadata. Table 1 provides a summary of the results. The complete row-level mapping table is available as a CSV file on Zenodo.[6]

Table 1: Coverage of GEO fields by the MeSyTo Metadata Model (MMM) and their occurrence in the retrieved NCBI metadata.

| **Category** | **Fields** | **Percentage** | **Denominator** |
|---|---|---|---|
| GEO fields included in the analysis | 32 | 100% | 32 GEO fields |
| Mapped to the MMM | 28 | 87.5% | 32 GEO fields |
| Also found in the NCBI metadata | 16 | 57.1% | 28 mapped fields |
| Not found in the NCBI metadata | 12 | 42.9% | 28 mapped fields |
| No MMM-specific mapping | 4 | 12.5% | 32 GEO fields |

[5] `https://codebase.helmholtz.cloud/department-computational-biology/software/mesyto`

[6] `https://zenodo.org/records/21931025/files/geo_mesyto_ncbi_crosswalk.csv`

Overall, the case study shows that the MMM can harmonize most fields in the GEO sequencing submission structure and apply additional validation rules needed for toxicological omics metadata.

### References

Canzler, S. *et al.* (2025). Evaluating the performance of multi-omics integration: a thyroid toxicity case study. *Arch Toxicol*, **99**(1), 309–332.
